# Whole-genome Studies of Malagasy People Uncover Novel Body Composition Associations

**DOI:** 10.1101/2023.11.21.568192

**Authors:** Iman Hamid, Séverine Nantenaina Stéphie Raveloson, Germain Jules Spiral, Soanorolalao Ravelonjanahary, Brigitte Marie Raharivololona, José Mahenina Randria, Mosa Zafimaro, Tsiorimanitra Aimée Randriambola, Rota Mamimbahiny Andriantsoa, Tojo Julio Andriamahefa, Bodonomena Fitahiana Laza Rafidison, Mehreen Mughal, Anne-Katrin Emde, Melissa Hendershott, Sarah LeBaron von Baeyer, Kaja A. Wasik, Jean Freddy Ranaivoarisoa, Laura Yerges-Armstrong, Stephane E. Castel, Rindra Rakotoarivony

## Abstract

The majority of human genomic research studies have been conducted in European ancestry cohorts, at the loss of detecting potentially novel and globally impactful findings. Here, we present the first whole genome sequence data and genome-wide association study in a cohort of 264 Malagasy individuals from three locations on the island of Madagascar. We describe genetic variation in this Malagasy cohort, providing insight into the shared and unique patterns of genetic variation across the island. We observe phenotypic variation by location, and find high rates of hypertension particularly in the Southern Highlands as well as elevated malaria prevalence in the West Coast relative to other sites. We find a number of genetic associations with body composition traits, including many variants that are unique to African populations or populations with admixed African ancestry such as Madagascar. This study highlights the utility of including diverse populations in genomic research for the potential to gain novel insights, even with small cohort sizes. This project was conducted in partnership and consultation with local Malagasy stakeholders and serves as an example for equitable genomic research with potential impacts on our understanding of human health and disease.

## Main Text

Humans have inhabited Madagascar for approximately 2000 years.^1–3^ The Malagasy population can trace their genetic ancestry to eastern African Bantu-speaking populations and Austronesian groups most likely from modern day Borneo, Indonesia.^4,5^ Austronesian founders arrived in eastern Madagascar and African founders in the northwest coast of Madagascar. These ancestral groups likely settled the island at different times and in waves over the course of 1000-2000 years, with subsequent admixture beginning approximately 28 generations in the past. Following a period of population structuring by geography there have been more recent internal migrations as a result of increased travel across regions within Madagascar.^6–9^

There have been few genome-wide studies of the Malagasy population of Madagascar. Of these existing studies, most have primarily focused on the selective and demographic history including the timing of settlement, evidence of past bottlenecks, admixture, and positive selection.^4,5,8,10,11^ The unique genetic architecture of Madagascar is of particular interest from a population genetics perspective. There is a known signal of strong post-admixture selection at the malaria protective Duffy-null variant on chromosome 1.^10,12^ Beyond these two studies which both focused on the single locus signal at Duffy-null, other signals of selection in Malagasy have not been investigated. Further, these past studies either focused on a single region in Madagascar (central highlands)^12^, or grouped individuals from multiple regions across the island into one population for analysis purposes.^10^ However, due to the settlement history of the island, patterns of genetic diversity vary across Madagascar, including relative contributions of Austronesian and African sources.^8^ Accordingly, there may be differences in selective pressures and genomic regions under selection for groups residing in different parts of Madagascar. The recent admixture and selective pressures unique to the island may have led novel or globally rare variants with functional impacts to increase in frequency. This provides a distinctive opportunity to detect genetic associations either at novel loci or with novel variants at known loci, which can add to our understanding of the underlying biology driving an association.

Of note, the genome-wide studies to date have all relied on genotyping array data. Here, we describe the first whole genome sequence data from the Malagasy population of Madagascar. We also identify several genetic associations with anthropometric phenotypes, representing the first GWAS in a Malagasy cohort. This study expands the diversity of populations that have been included in genomic research, allowing us to gain novel insight into human health both in Madagascar and globally.

This study was conducted in collaboration with scientists and anthropologists at Variant Bio (VB) and the University of Antananarivo Department of Anthropobiology and Sustainable Development (UA). First, representatives of VB and UA conducted an initial community consultation in order to determine study sites and to assess the feasibility and interest in genetic research within the community. We also established a benefit-sharing agreement based on the needs of the community, with access to clean water and improved school infrastructure being the most common concerns across communities.13

Ultimately, the study included community-based recruitment of adult volunteers from three locales in order to gain population-representative samples: Tsianaloka Village, Tsimafana (West Coast); Ampandrialaza Village, Ampangabe (Central Highlands); and Tsiandatsiana Village, Ankerana (Southern Highlands) (Figure 1A). We collected 15 anthropometric and spirometric phenotypes as well as history of malaria infection. See Supplemental Methods for more details on participant inclusion criteria and phenotyping.

**Figure 1.**
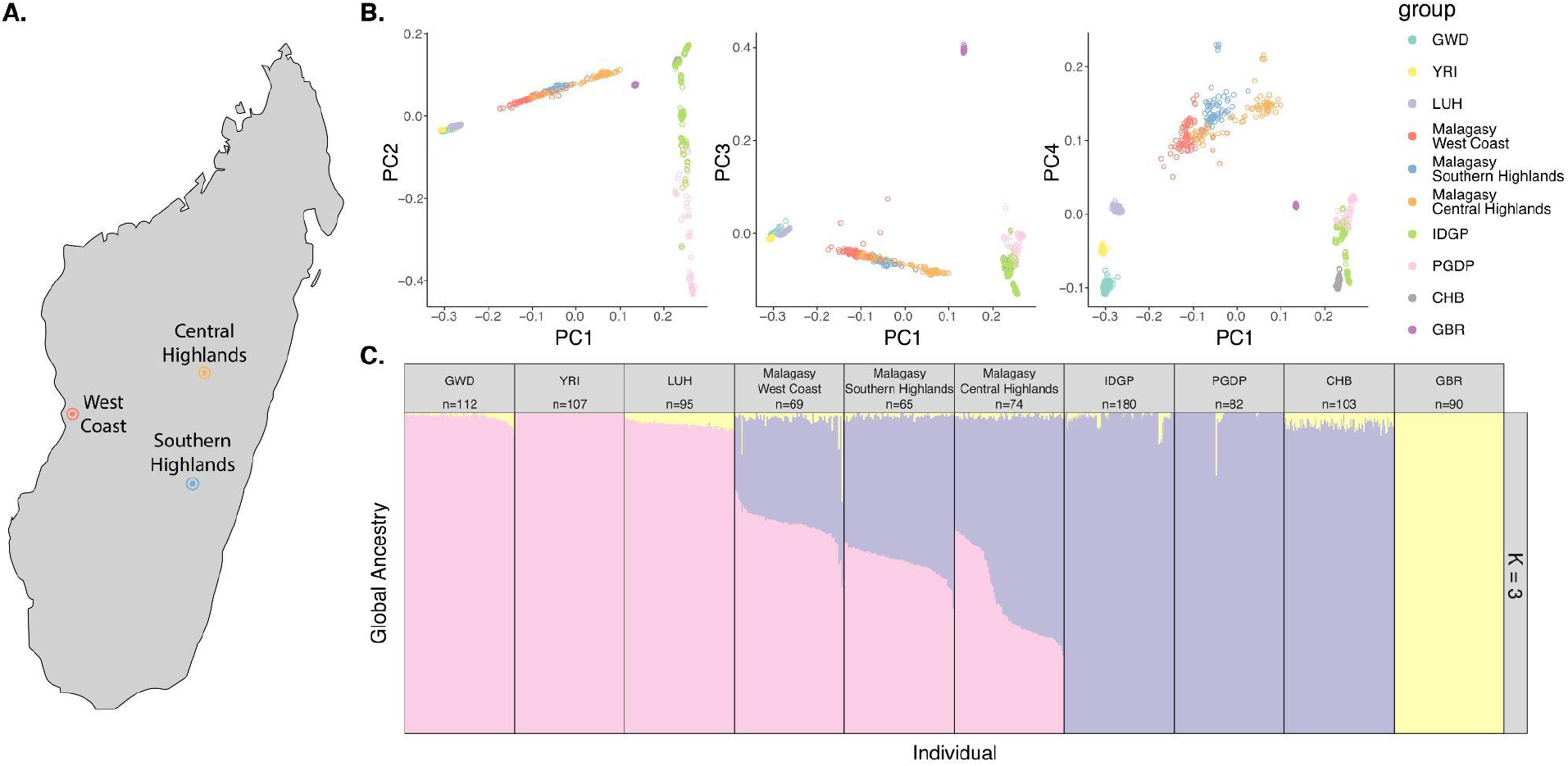
Cohort overview. **A)** Map of three sampling sites on Madagascar corresponding to the West Coast (Tsianaloka Village, Tsimafana), Central Highlands (Ampandrialaza Village, Ampangabe), and the Southern Highlands (Tsiandatsiana Village, Ankerana). **B)** PCs 2-4 vs PC1 and **C)** ADMIXTURE plot for K=3, including Malagasy cohort (West Coast, Central Highlands, Southern Highlands) and relevant reference populations from 1000 Genomes (GWD, YRI, LUH CHB, GBR) and additional reference samples from IDGP and PGDP. Points in the scatterplot are colored by group legend, global ancestry bars are colored by proportion of African-related (pink), Asian-related (purple), or European-related (yellow) ancestry.

Overall, the participants were healthy, with the vast majority of individuals self-reporting excellent health (Figure 2A). The sample population was generally young, with an average age across the three sites ranging from 29 to 35 years old (West Coast = 35 years, Southern Highlands = 33 years, Central Highlands = 29.5 years) (Figure 2B, Supplemental Table 1). Approximately 20% of participants had been to a doctor in the year preceding this study. Only 14 individuals (∼5%) reported having malaria in the past year, and 20% of participants reported ever having malaria. This proportion of both male and female participants reporting ever having malaria is noticeably higher in the West Coast (male = 56%, female = 41%) compared to the other two study sites (0-11%), likely reflective of an increased presence of the malaria parasites along the coast (Supplemental Table 2). We measured both systolic and diastolic blood pressure and categorized individuals who had Stage 1 or Stage 2 hypertension versus normal blood pressure based on American College of Cardiology (ACC) guidelines.14 We find that approximately 61% of participants were hypertensive (Normotensive / slightly elevated = 39%, Stage 1 hypertension = 31% Stage 2 hypertension = 30%), with the Southern Highlands showing on average higher blood pressure than the other two sites (Figure 2C, Supplemental Table 1). A previous study has similarly reported high rates of hypertension (49%) in a rural community in the northeastern coast of Madagascar.15 In Figure 2C, we show distributions of six phenotypes stratified by site and sex: maximum forced expiratory volume in 1 second (FEV1 max) measured in percentage, systolic blood pressure measured in millimeters of mercury (mmHg), weight measured in kilograms (kg), body fat percentage, and hip and weight circumferences measured in centimeters (cm). Descriptive statistics for all collected phenotype distributions, stratified by site and sex, are provided in Supplemental Tables 1 & 2.

**Figure 2.**
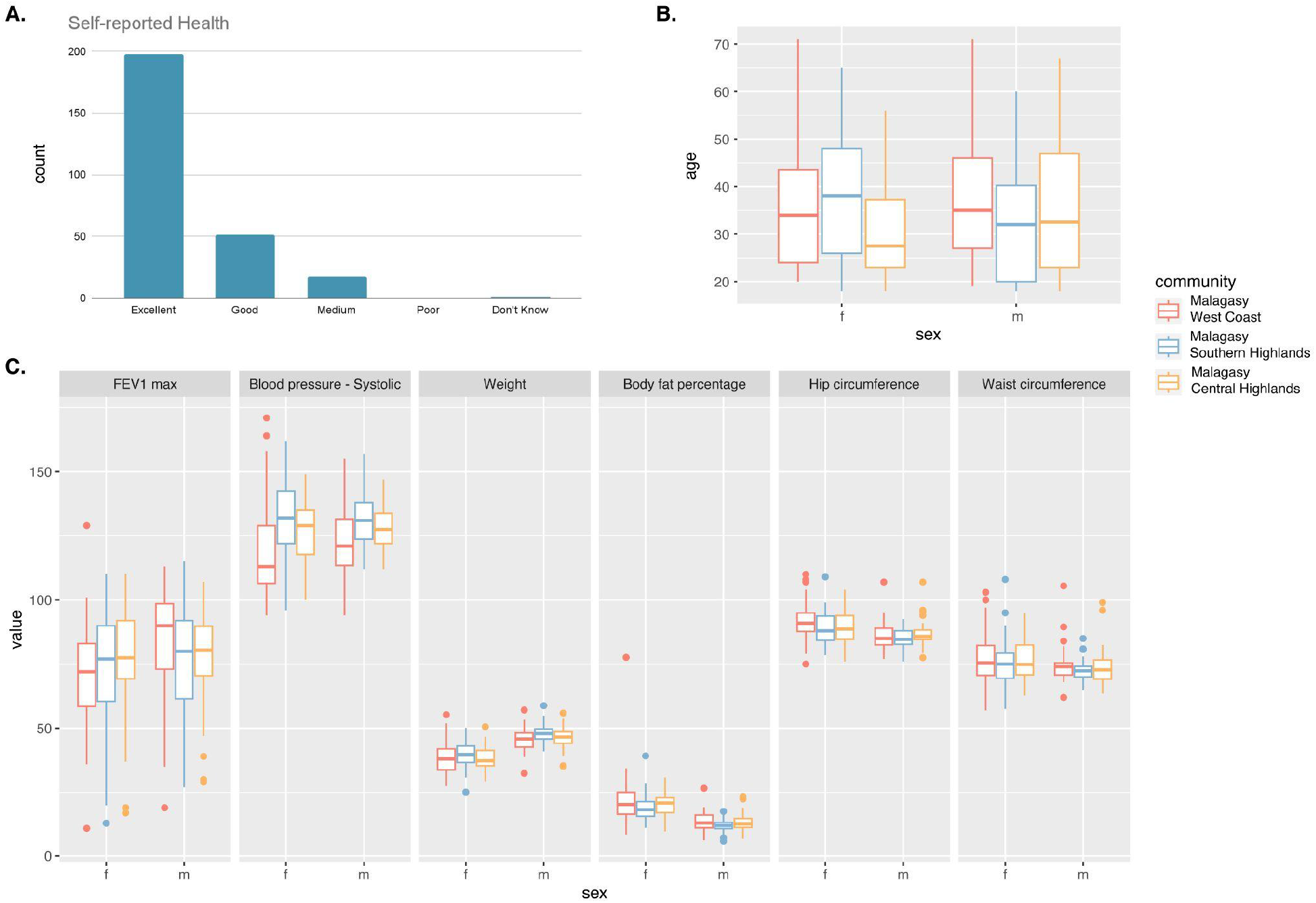
Phenotype distributions by site. **A)** Self-reported health across all three sites **B)** Distribution of ages stratified by site (color) and sex (x-axis) **C)** Distribution of traits values stratified by site and sex for six of the collected phenotypes (FEV1 max [percent], systolic blood pressure [mmhg]], weight [kg], body fat percentage [percent], hip circumference [cm], waist circumference [cm]).

Mid-pass whole genome sequencing, joint genotyping, and imputation was performed for all participants, with 264 passing quality control at a mean coverage of 4.6x (Supplemental Methods)16. Reference genomes were included from the Indonesian Genome Diversity Project (IDGP) (EGAS00001003054) and the Papuan Genome Diversity Project (PGDP)17 and 1000 Genomes references from eastern Asia, Europe, and central, eastern, and western Africa (Han Chinese [CHB], British [GBR], Yoruba [YRI], Luhya [LWK], Gambian [GWD]).18 Including the Malagasy cohort, the final call set included 1,077 individuals. A summary table of population-level allele frequencies for the ∼29 million alleles segregating in the Malagasy cohort is provided (see Data Availability).

We performed separate quality checks and filtering at the individual and variant level for GWAS and population genetics analyses respectively (Supplemental Methods). We ran a principal components analysis (PCA) in Hail (v0.2) and ADMIXTURE for a subset of 977 individuals passing QC, including 208 of the Malagasy participants.19,20 Individuals within the Malagasy cohort cluster by locale along the African to Asian ancestry cline in PC space (Figure 1B). The clustering by site is likely indicative of different proportions of African and Austronesian ancestry resulting from the complex settlement history of the island. Indeed, we see from ADMIXTURE results at K=3 that the average proportion of African-related global ancestry is higher in the West Coast (66% African) compared to a higher average proportion of Austronesian-related global ancestry in the Central Highlands (41% African), with the global ancestry proportions Southern Highlands being closer to equal contributions from the two ancestral sources (53% African) (Figure 1C).

We identified genetic variants that are unique or enriched in the Malagasy population relative to other reference populations. Specifically, we defined a variant as “enriched” if the minor allele frequency was less than 0.01 in gnomAD v3,21,22 less than 0.01 in all reference populations included in joint calling and imputation (LUH, GAM, YRI, CHB, GBR, IDGP, PGDP) and more than 0.05 in the Malagasy cohort. We further annotated these enriched variants as having likely functional impacts if they had a CADD greater than 30 or else was annotated by Variant Effect Predictor with a consequence of “high” or “moderate” impact according to the Ensembl IMPACT rating (https://ensembl.org/info/genome/variation/prediction/predicted_data.html) (e.g. stop gain, stop lost, frameshift, missense, etc). Specifically, “high” impact variants are defined as those that are “assumed to have high (disruptive) impact in the protein, probably causing protein truncation, loss of function or triggering nonsense mediated decay,” and “moderate” impact variants are defined as those that are “non-disruptive” and “might change protein effectiveness.” Out of 44,731,585 total variants passing QC, we identified 20,656 variants that were enriched in the Malagasy population (maf > 0.05) and either absent or low frequency (<0.01) in other global reference populations as described above. Of these, 116 had likely functional impact according to our criteria (Figure 3A, Supplementary File 1).

**Figure 3.**
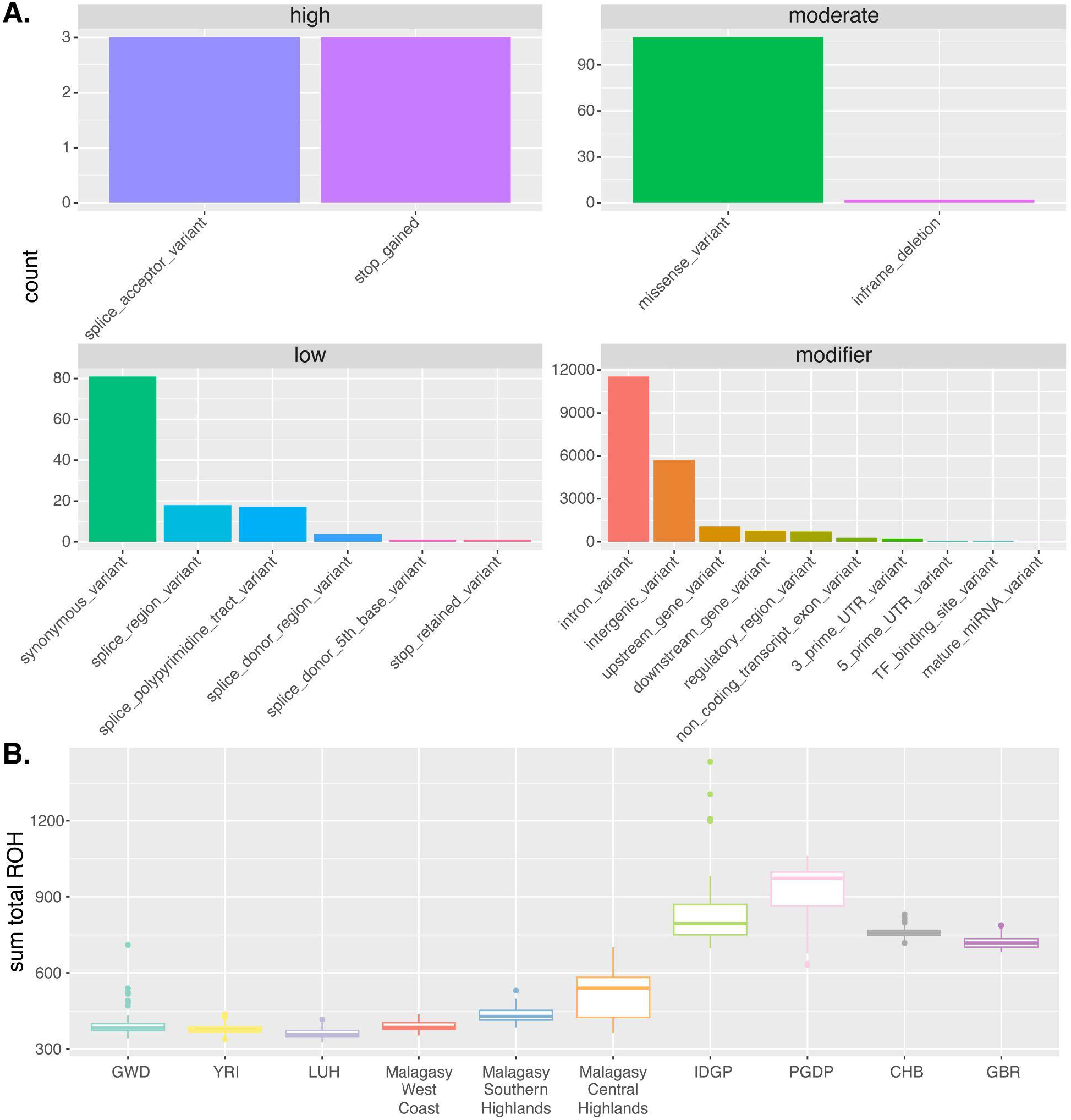
Genetic Variation Overview. **A)** Count of variants enriched in the Malagasy cohort relative to global populations, grouped by functional category (VEP consequence, x-axis) and Ensembl IMPACT classification (high, medium, low, moderate). **B)** Sum of the genome contained in runs of homozygosity (ROH) by Madagascar study site or reference population.

Next, runs of homozygosity (ROH) were called to identify signatures of recent or older bottlenecks. We detected runs of homozygosity (ROH) for each individual using bcftools/RoH.23 We called ROH for each individual, and calculated the total amount of the genome contained within ROH. Larger amounts of the genome contained in ROH, and in particular ROH spanning long genomic distances, may be indicative of recent bottleneck events.24 In Figure 3B, we see that the overall patterns of ROH align with expectations. That is, bottlenecked out-of-Africa (OOA) populations showing higher sum total amounts of the genome contained in ROHs and continental African populations show lower values due to the higher effective population sizes. Among the three study sites in Madagascar, the Central Highlands have higher values of sum total ROH, reflective of a historical bottleneck in the region. This is consistent with a past study which found evidence of a bottleneck in a central highlands subgroup based on patterns of IBD sharing from SNP array data.8,25

We performed GWAS for the 16 phenotypes collected across 12,528,464 million variants passing GWAS QC (Supplemental Methods), combining all individuals from across the three sampling sites into one cohort due to the low sample size (213-214 individuals, varying by phenotype). We identified 22 variants reaching genome-wide significance (p < 5e-8) for at least one phenotype. Summary statistics for the lead variant associations are shown in Table 1, variant-level summary statistics for all significant associations can be found in Supplemental Table 3, and an additional summary of all GWAS is provided in Supplemental Table 4. Full summary statistics are provided for all GWAS (Data Availability).

**Table 1.**
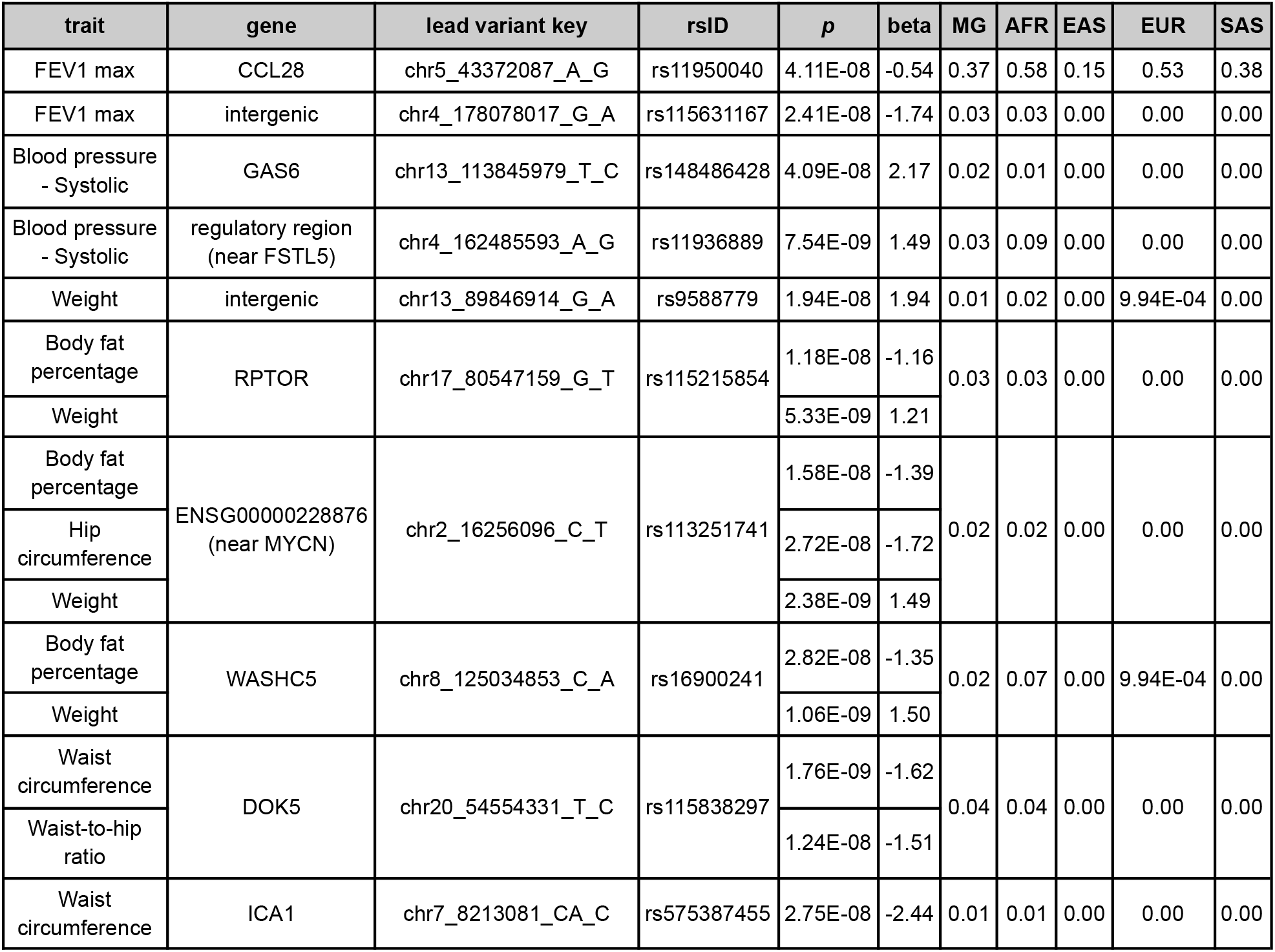
Summary statistics for the lead variants from 15 significant associations. The *p*-values and beta coefficients for each lead variant and trait associated locus are listed. The lead variant key (chr_pos_ref_alt), rsID, VEP annotated gene (or closest coding gene in parenthesis), and allele frequencies are included (MG=Malagasy cohort from this study, AFR=1000 Genomes African populations, EAS=1000 Genomes East Asian populations, EUR=1000 Genomes European populations, SAS=1000 Genomes South Asian populations).

| trait | gene | lead variant key | rsID | *p* | beta | MG | AFR | EAS | EUR | SAS |
| --- | --- | --- | --- | --- | --- | --- | --- | --- | --- | --- |
| FEV1 max | CCL28 | chr5_43372087_A_G | rs11950040 | 4.11E-08 | -0.54 | 0.37 | 0.58 | 0.15 | 0.53 | 0.38 |
| FEV1 max | intergenic | chr4_178078017_G_A | rs115631167 | 2.41E-08 | -1.74 | 0.03 | 0.03 | 0.00 | 0.00 | 0.00 |
| Blood pressure<br>- Systolic | GAS6 | chr13_113845979_T_C | rs148486428 | 4.09E-08 | 2.17 | 0.02 | 0.01 | 0.00 | 0.00 | 0.00 |
| Blood pressure<br>- Systolic | regulatory region<br>(near FSTL5) | chr4_162485593_A_G | rs11936889 | 7.54E-09 | 1.49 | 0.03 | 0.09 | 0.00 | 0.00 | 0.00 |
| Weight | intergenic | chr13_89846914_G_A | rs9588779 | 1.94E-08 | 1.94 | 0.01 | 0.02 | 0.00 | 9.94E-04 | 0.00 |
| Body fat<br>percentage | RPTOR | chr17_80547159_G_T | rs115215854 | 1.18E-08 | -1.16 | 0.03 | 0.03 | 0.00 | 0.00 | 0.00 |
| Weight |  |  |  | 5.33E-09 | 1.21 |  |  |  |  |  |
| Body fat<br>percentage | ENSG00000228876<br>(near MYCN) | chr2_16256096_C_T | rs113251741 | 1.58E-08 | -1.39 | 0.02 | 0.02 | 0.00 | 0.00 | 0.00 |
| Hip<br>circumference |  |  |  | 2.72E-08 | -1.72 |  |  |  |  |  |
| Weight |  |  |  | 2.38E-09 | 1.49 |  |  |  |  |  |
| Body fat<br>percentage | WASHC5 | chr8_125034853_C_A | rs16900241 | 2.82E-08 | -1.35 | 0.02 | 0.07 | 0.00 | 9.94E-04 | 0.00 |
| Weight |  |  |  | 1.06E-09 | 1.50 |  |  |  |  |  |
| Waist<br>circumference | DOK5 | chr20_54554331_T_C | rs115838297 | 1.76E-09 | -1.62 | 0.04 | 0.04 | 0.00 | 0.00 | 0.00 |
| Waist-to-hip<br>ratio |  |  |  | 1.24E-08 | -1.51 |  |  |  |  |  |
| Waist<br>circumference | ICA1 | chr7_8213081_CA_C | rs575387455 | 2.75E-08 | -2.44 | 0.01 | 0.01 | 0.00 | 0.00 | 0.00 |

Our results include many low to moderate frequency variants specific to African ancestry populations associated with body composition traits (Table 1, Supplemental Table 3). For example, a haplotype overlapping with RPTOR on chromosome 17 was associated with both body fat percentage and weight (Figure 4A, Table 1). The lead variant (rs115215854) is an intron variant in RPTOR. While variants in RPTOR have previously been associated with body mass index (BMI) in biobank-scale studies (UKBiobank, FinnGen), these cohorts had sample sizes on the order of 450-800k individuals.26–29 With a fraction of that sample size (n = 214), we detect significant associations at RPTOR. Further, the lead variant for these two body composition associations is segregating at low frequencies (∼ 3%) in African populations or populations with admixed African ancestry, including the Malagasy cohort, and unobserved in other 1000Gs populations around the world (Table 1).

**Figure 4.**
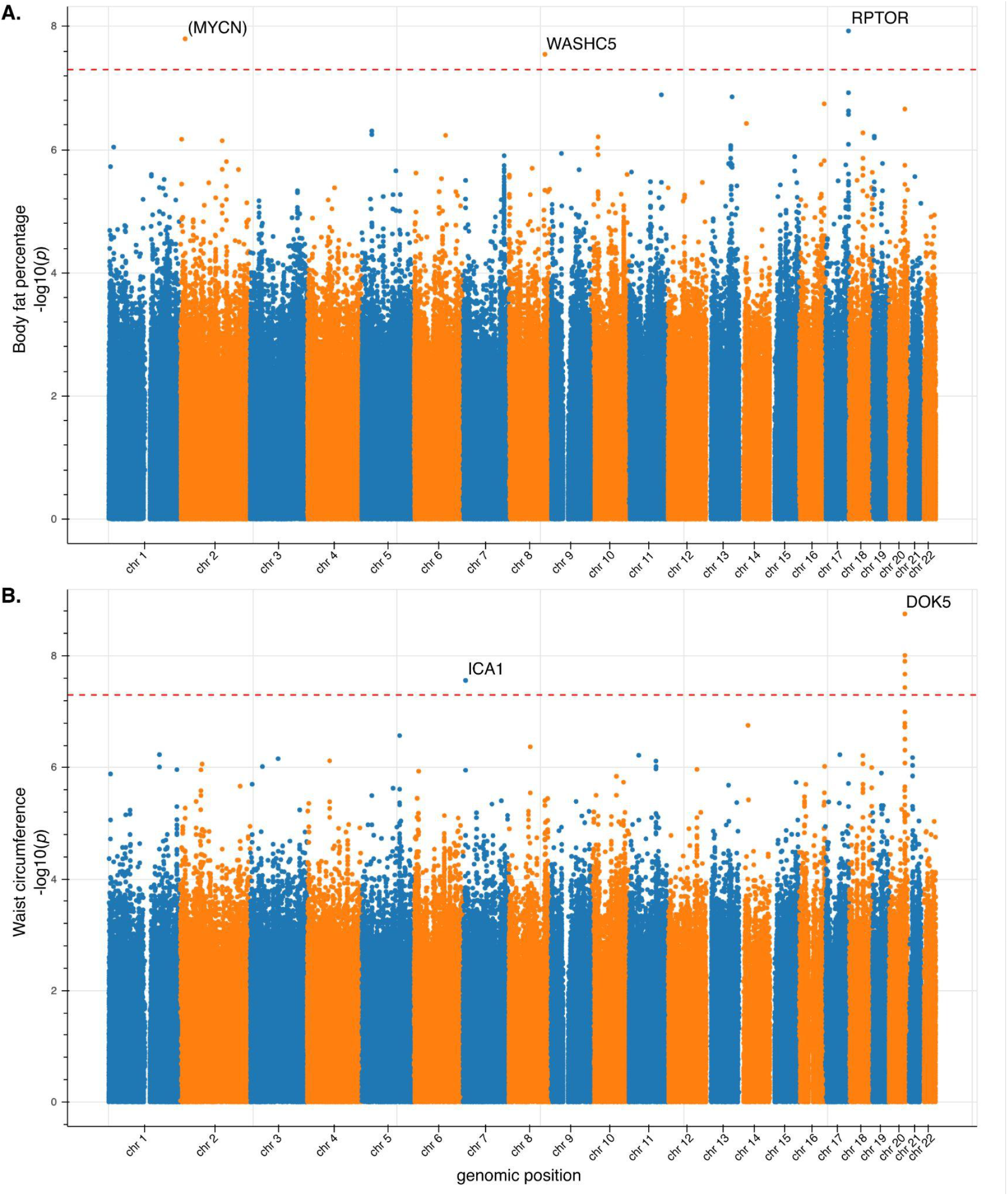
GWAS Manhattan plots for two anthropometric traits. *P*-values for all autosomal variants passing GWAS QC on a −log10 scale for **A)** body fat percentage and **B)** waist circumference. Annotated gene for top variant, or nearest protein coding gene in parenthesis, is listed for each locus reaching genome-wide significance (*p* < 5e-8, red dashed line).

Similarly, we identified a significant association with waist circumference and waist-to-hip ratio on chromosome 20, with the lead variant (rs115838297) in the intron of DOK5 (Figure 4B). Variants in DOK5 have been weakly associated with body mass index in the UKBiobank with large cohort sizes.26,27,30 Again, we calculated the allele frequencies for the lead variant from this study in 1000 Genomes populations and found that this allele is unique to African populations or populations with admixed African ancestry, including the Malagasy cohort and segregates at a low frequency (∼ 4%) (Table 1).

Among the novel associations is a variant in WASHC5 on chromosome 20 that is positively associated with weight and negatively associated with body fat percentage (rs16900241) (Figure 4A). We find no past studies associating variants in WASHC5 with body composition traits. WASHC5 encodes the protein strumpellin which is most highly expressed in skeletal muscle and also highly expressed in the thyroid.31 We also identified a significant association with the maximum forced expiratory volume in 1 second (FEV1 max) on chromosome 5. The peak variant (rs11950040) is globally common with an allele frequency of 42.8% in 1000 Genomes, and is a downstream gene variant for CCL28 (Table 1, Supplemental Table 3). Additional summary statistics, including population-level allele frequencies for lead variants and all significant variant associations are provided in Table 1 and Supplemental Table 3, respectively.

Despite the small cohort size in this study, GWAS for well-powered quantitative body composition traits like weight and waist circumference led to discovery of novel associations or novel variants at known trait-associated loci. We included a minor allele count (MAC) threshold in our QC criteria (see Supplemental Methods); however, we recognize that this cohort is a small sample size and that a number of our GWAS associations are with variants between 1 and 5% minor allele frequency. Though we urge caution in interpretation, we are encouraged that a number of the low frequency variant associations are in genes that have previously been associated with relevant phenotypes in larger cohorts (e.g. DOK5, RPTOR). Additionally, the lead variant associated with FEV1 max, chr5_43372087_A_G / rs11950040, is also common in European ancestry and has a nominally significant association with peak expiratory flow in Europeans.32 Although we combined the cohort across sites due to low sample sizes, the slight genetic differentiation and varying environmental or selective pressures across the sites suggests deeper sampling and investigations in each region could identify group-specific genetic associations with traits of interest.

This study of genetic and phenotypic variation across three Malagasy villages demonstrates site-specific trait distributions including malaria incidence and genetic substructure by locale. These results emphasize the need for more granular genomic studies that take into account region-specific genetic and environmental differences. In particular, these results have implications for studies that have scanned the Malagasy genome for signals of positive selection. Studies have identified the strong selection at Duffy-null in a cohort with participants either grouped together from all across the island of Madagascar10 or at one specific site.12 Instead, a study designed to detect selection signals in each site independently may find that the post-admixture Duffy-null signal is not as wide-spread, particularly in places where malaria incidence is low. A site-specific study design may also find unique and novel selection signals at each locale that were masked in a study that grouped all populations despite the clear substructuring that we observe in our analysis. The WGS data described in this study from three specific sites is a valuable start for identifying such region-specific signals.

The Malagasy people have never been included in WGS studies or GWA studies, but here we show that even with a small cohort, we can draw potentially important insights into the genetic basis of clinically relevant traits. This study demonstrates the utility of partnering with local communities in order to expand the diversity of people who participate in and potentially benefit from genomic research.

## Supplemental Information

Supplemental Materials includes Supplemental Methods, four tables and four figures.

## Acknowledgments

We would like to thank all the Malagasy people who participated in the study. We acknowledge Anthropobiologie et Développement Durable, Ecole Doctorale Sciences de la Vie et de l’Environnement, Université d’Antananarivo and all participating communities and in-field contributors. We also thank Leslie Hepner and Noah Collins for their assistance with training, translation, and study execution.

## Ethics Approval and Consent to Participate

Participants were required to be at least 18 years old, not pregnant, and willing and able to consent. Participants signed an informed consent form approved by the ethics committee Comité Malgache d’Ethique Pour les Sciences et les Technologies (CMEST).

## Data and Code Availability

Population-level allele frequencies for all alleles segregating in the Malagasy cohort (i.e. variants with alternate allele frequencies between 0 and 1 in Madagascar) are available at https://public.variantbio.com/MGUA. This summary table includes reference population allele frequencies from 1000G, gnomAD, IDGP, PGDP and relevant variant annotations such as CADD, VEP annotated gene, and predicted functional consequence. Annotated variant-level summary statistics are provided for all variants that were included in GWAS. Full GWAS summary statistics were uploaded to the GWAS Catalog for each of the 16 phenotypes (accession numbers: GCST90297828, GCST90297829, GCST90297830, GCST90297831, GCST90297832, GCST90297833, GCST90297834, GCST90297835, GCST90297836, GCST90297837, GCST90297838, GCST90297839, GCST90297840, GCST90297841, GCST90297842, GCST90297843). Detailed workflow and scripts for mid-pass WGS variant calling and imputation are available on GitHub: https://github.com/variant-bio/mid-pass.

## Author Contributions

I.H. and S.N.S.R. analyzed data and wrote the manuscript. R.R., G.J.S., S.R., J.F.R., B.M.R., J.M.R., K.A.W., S.L.V.B., L.Y.A. and S.E.C. designed the study. S.N.S.R., R.R., M.H., G.J.S., S.R., J.F.R., B.M.R., J.M.R., M.Z., T.A.R., R.M.A., T.J.A., B.F.L.R. and S.L.V.B. carried out the study. S.N.S.R., R.R., G.J.S., S.R., J.F.R., B.M.R., J.M.R., M.Z., T.A.R., R.M.A., T.J.A., B.F.L.R., and S.E.C. processed and quality controlled all subject phenotype and meta data. A.E. processed sequencing data, performed imputation, and quality control. M.M. carried out population genetic analyses. R.R. and S.L.V.B. performed community engagement and consultation. S.E.C. carried out GWAS. All authors reviewed and contributed to writing the manuscript.

## Supplementary Materials

### Supplemental Methods

#### Study design and sampling protocol

This study was designed in collaboration between Variant Bio and the University of Antananarivo Department of Anthropobiology and Sustainable Development. The first phase of study design consisted of three weeks of formal community engagement in three locations across Madagascar. The research team employed semi-structured interviews with community leaders, key stakeholders, and other community members both to explain the purpose of the project, assess local interest and study feasibility, and discuss their questions and concerns, as well as to gather input from community members about how best to allot benefit sharing funds.

Preliminary questions about family and ancestry were asked, and we required self-reported ancestry from all four grandparents to be from the same region of Madagascar. Participants were asked to refrain from food and drink within the 30 minutes of sample collection. DNA was extracted from saliva samples using the DNA Genotek Oragene kit.

#### Phenotyping

Anthropometrics were measured for each participant, including height, weight, hip and waist circumference, muscle mass, body fat (bioimpedance scale), biacromial width, biiliac width, grip strength, arm circumference, triceps skinfold measurement, blood pressure, and body temperature. For blood pressure, hypertension was defined using the American College of Cardiology (ACC) guidelines (Whelton et al. 2018). Specifically, Stage 1 hypertension was defined as systolic blood pressure from 130 to 139 or diastolic blood pressure from 80 to 89 mm Hg and Stage 2 hypertension was defined as systolic blood pressure greater than or equal to 140 mm Hg or diastolic blood pressure greater than or equal to 90 mm Hg. A spirometer was used to calculate the maximum forced expiratory volume in one minute (FEV1) and the maximum forced vital capacity (FVC max) across three measurements. Data were de-identified prior to analysis.

#### WGS and data processing

##### Sequencing, sample processing and QC

Saliva samples from 266 individuals were sent to Psomagen, Inc. (Maryland, USA) for DNA extraction, library preparation and sequencing. DNA samples were prepared with Illumina DNA Prep library kits and sequenced on NovaSeq 6000 instruments with 2×151bp reads targeting throughput of ∼16Gb/sample, yielding mean deduplicated coverage of 4.6x per sample on average.

Processing was done with Variant Bio’s in-house processing pipeline. Raw sequencing data were inspected with fastqc (v0.11.7) and adapters were trimmed using cutadapt (v2.10). Trimmed reads were then processed following the GATK Best Practices guidelines, more specifically the CCDG functional equivalence version (Regier et al. 2018) (using BWA-mem v0.7.15 for mapping onto GRCh38, Picard v2.21.4 for duplicate marking, samtools v1.10 for sorting, and GATK v4.1.4.0 for BQSR). Coverage was determined using Picard CollectWgsMetrics. Two samples were found to have excessive bacterial contamination with mapping rates to GRCh38 below 30% and excluded from further analysis.

##### Variant calling and imputation

Variant calling was performed using HaplotypeCaller (GATK 4.1.4.0) in GVCF mode, specifying the parameter “--read-filter OverclippedReadFilter” to disregard short alignments (largely from bacterial contamination). Joint genotyping was performed using genomicsDBImport and GenotypeGVCFs (GATK 4.2.0.0) together with 514 individuals from the 1000 Genomes Project (Han Chinese [CHB], British [GBR], Yoruba [YRI], Luhya [LWK], Gambian [GWD]; all sequenced at high coverage and processed with the same pipeline). We next ran VQSR (setting --truth-sensitivity-filter-level to 99.8 for SNPs and 99.0 for indels) and only retained PASS filter sites for further analyses. Next, we ran imputation within the cohort using Beagle v5.1 (beagle.27Apr20.b81.jar), with all genotypes with GQ < 18 set to missing (Emde et al. 2021) (method: https://github.com/variant-bio/mid-pass). In a first pass, we only imputed genotypes of variants in Genome In a Bottle (GIAB) “easy” regions (Olson et al. 2022), ensuring high quality imputation within this stricter set. After fixing these genotypes, we added variants outside easy regions as well as structural variants (SVs) from the Human Genome Structural Variation Consortium genotyped in all individuals with PanGenie (Ebler et al. 2022) (GQ < 200 set to missing) and imputed these within the cohort as well.

Imputed genotypes were loaded into Hail v0.2 and annotated with VEP (v108) and CADD Phred (SNPs only, v1.6), sub-population specific allele frequencies, as well as impute rate (defined as the proportion of genotypes that have been imputed, i.e. were missing or had GQ < 18).

##### Population genetic analysis QC & Subject / Variant Filtering

For population genetic analyses such as PCA, ADMIXTURE, and runs of homozygosity (ROH), we started with the joint callset which included 44,731,585 and 1,077 reference and Malagasy cohort samples. We first masked GIAB low confidence regions (Olson et al. 2022), resulting in 28,670,287 variants. We filtered 11 subjects with missing sex, 60 subjects with relatedness above 3rd degree (pihat > 0.125), and 29 subjects greater than 7 standard deviations from the mean for any of PCs 1-10 (in the unfiltered callset). This resulted in 977 total subjects, including 208 from the Malagasy cohort (West Coast n = 69; Southern Highlands n = 65; Central Highlands n = 74). We filtered 740,169 variants with MSQ ranksum not equal to zero or variants that did not pass QC in gnomAD v3 (Karczewski et al. 2020). We filtered 140,648 variants without a MAC of at least 5 after subject-level filtering. We again restricted the analyses to autosomes only, resulting in an additional 907,092 variants filtered. Ultimately, this final callset included 977 subjects and 26,882,378 variants.

PCA was run in Hail v0.2 with this filtered call set using LD pruned variants (MAF > 0.01, *r*^2^ < 0.20) and the “hwe_normalized_pca” function with “k = 20”. Prior to running ADMIXTURE and identifying runs of homozygosity (ROH), variants were additionally filtered to include only biallelic SNPs with a minor allele frequency (MAF) greater than 0.05. We identified runs of homozygosity (ROH) from this VCF for each individual using bcftools/RoH (Narasimhan et al. 2016). This set was further pruned for LD (*r*^2^ < 0.4 in 50,000 bp windows) and then randomly downsampled to include 150,000 total variants genome-wide. ADMIXTURE was run unsupervised for K=2 to K=5 on the LD-pruned and downsampled VCF (Alexander et al. 2009).

#### Genome-wide association studies

##### GWAS Subject / Variant Filtering

For GWAS, the joint callset, which included 1077 reference and Malagascy cohort samples, was filtered to include only those subjects from the Malagasy cohort (n = 264). Six subjects were filtered due to missing sex. One subject was excluded for heterozygosity greater than three standard deviations from the population mean. 42 subjects were filtered due to relatedness greater than fourth degree (pihat > 0.0625). The final cohort for GWAS included 214 Malagasy individuals (West Coast n=79; Southern Highlands n=67; Central Highlands n=68). Of the 44,731,585 variants in the joint callset, we filtered 24,689,620 variants without a minor allele count (MAC) of at least 5 (corresponding to an allele frequency of 1.17%) after subject-level filtering and 4,536,292 indels in low-confidence regions or variants that did not pass QC in gnomAD v3 (Karczewski et al. 2020). We used an iterative variant level call rate / impute rate filtering with iterative rates of 0.85, 0.8, and 0.75, which filtered an additional 2,833,411 variants. We removed 60,923 variants with *p* = 0 for the HWE test. We restricted the analyses to autosomes only, resulting in an additional 82,875 variants filtered. Ultimately, the final set included 214 subjects and 12,528,464 variants that passed QC for GWAS.

##### GWAS

All quantitative traits were inverse-normal transformed prior to running GWAS. The quality controlled and filtered callset was used to run GWAS in Hail v0.2 with the “linear_regression_rows” function for quantitative traits and the “logistic_regression_rows” function for binary traits. The following covariates were used in the regressions: sex + age + age^2^ + sex*age + sex*age^2^ + mean depth + genetic principal components 1-20. Principal components were calculated using the same subjects as the GWAS. Genomic inflation (λ GC) was calculated for each GWAS using the “lambda_gc” function with “approximate = False”. Genome-wide significance for each GWAS was defined as p < 5×10^−8^.

## Supplemental Figures and Tables

**Supplemental Table 1.** Descriptive statistics for continuous phenotypes collected stratified by site and community. For each quantitative trait we provide the range [min, max], mean, standard deviation, 95% confidence interval, median, quartile 1, quartile 3 [Q1,Q3], and total number of samples with a value for each phenotype.

| phenotype | site | sex | range | mean | sd | CI (95%) | median | [Q1,Q3] | n |
| --- | --- | --- | --- | --- | --- | --- | --- | --- | --- |
| Age (years) | Central Highlands | f | [18,56] | 31.23 | 10.14 | [28.14,34.31] | 27.50 | [23.00,37.25] | 44 |
|  |  | m | [18,67] | 35.28 | 12.92 | [31.45,39.12] | 32.50 | [23.25,46.50] | 46 |
|  | Southern Highlands | f | [18,65] | 36.88 | 13.42 | [32.75,41.01] | 38.00 | [26.50,47.00] | 43 |
|  |  | m | [18,60] | 32.11 | 12.41 | [28.34,35.89] | 32.00 | [20.00,40.25] | 44 |
|  | West Coast | f | [20,71] | 36.05 | 13.97 | [31.80,40.29] | 34.00 | [24.00,43.50] | 44 |
|  |  | m | [19,71] | 36.72 | 12.52 | [32.87,40.57] | 35.00 | [27.50,45.50] | 43 |
| FEV1 max (%) | Central Highlands | f | [17.00,110.00] | 75.16 | 21.14 | [68.73,81.59] | 77.50 | [69.25,92.00] | 44 |
|  |  | m | [29.00,107.00] | 76.93 | 18.57 | [71.42,82.45] | 80.50 | [70.50,89.75] | 46 |
|  | Southern Highlands | f | [13.00,110.00] | 70.84 | 24.42 | [63.32,78.35] | 77.00 | [60.50,90.00] | 43 |
|  |  | m | [27.00,115.00] | 75.84 | 22.91 | [68.88,82.81] | 80.00 | [61.50,92.00] | 44 |
|  | West Coast | f | [11.00,129.00] | 69.59 | 21.15 | [63.16,76.02] | 72.00 | [58.50,83.00] | 44 |
|  |  | m | [19.00,113.00] | 83.28 | 21.25 | [76.74,89.82] | 90.00 | [73.00,98.50] | 43 |
| FVC max (%) | Central Highlands | f | [18.00,112.00] | 78.34 | 19.28 | [72.48,84.20] | 83.50 | [73.50,89.00] | 44 |
|  |  | m | [32.00,107.00] | 81.15 | 15.25 | [76.62,85.68] | 82.50 | [73.00,91.50] | 46 |
|  | Southern Highlands | f | [19.00,118.00] | 75.95 | 23.27 | [68.79,83.11] | 82.00 | [65.00,92.00] | 43 |
|  |  | m | [33.00,117.00] | 80.93 | 18.97 | [75.16,86.70] | 86.50 | [69.25,94.25] | 44 |
|  | West Coast | f | [10.00,118.00] | 71.82 | 19.85 | [65.78,77.85] | 73.00 | [63.00,82.25] | 44 |
|  |  | m | [32.00,116.00] | 83.65 | 16.56 | [78.56,88.75] | 89.00 | [75.00,95.00] | 43 |
| Height (cm) | Central Highlands | f | [141.40,167.40] | 153.22 | 6.62 | [151.21,155.23] | 152.75 | [148.65,156.98] | 44 |
|  |  | m | [151.80,180.00] | 164.60 | 6.16 | [162.77,166.43] | 164.60 | [161.70,168.78] | 46 |
|  | Southern Highlands | f | [142.20,167.60] | 152.47 | 5.63 | [150.73,154.20] | 152.20 | [148.50,155.90] | 43 |
|  |  | m | [152.90,177.40] | 163.75 | 5.84 | [161.97,165.53] | 162.95 | [159.13,168.08] | 44 |
|  | West Coast | f | [139.00,164.30] | 153.71 | 5.71 | [151.98,155.45] | 153.50 | [150.00,157.73] | 44 |
|  |  | m | [156.90,179.00] | 165.79 | 4.91 | [164.28,167.30] | 165.20 | [162.75,169.20] | 43 |
| Weight (kg) | Central Highlands | f | [29.30,50.60] | 38.15 | 4.64 | [36.74,39.56] | 37.30 | [35.35,41.35] | 44 |
|  |  | m | [35.10,55.90] | 46.17 | 4.27 | [44.90,47.43] | 46.55 | [44.13,48.60] | 46 |
|  | Southern Highlands | f | [25.10,50.10] | 39.71 | 5.28 | [38.09,41.34] | 39.80 | [36.65,43.10] | 43 |
|  |  | m | [41.00,58.80] | 47.68 | 3.78 | [46.53,48.82] | 47.90 | [45.80,49.75] | 44 |
|  | West Coast | f | [27.50,55.30] | 38.19 | 6.35 | [36.26,40.12] | 38.05 | [33.80,41.93] | 44 |
|  |  | m | [32.40,57.10] | 45.54 | 4.61 | [44.12,46.96] | 45.80 | [42.65,48.15] | 43 |
| Lean body mass (kg) | Central Highlands | f | [37.40,71.70] | 53.30 | 7.88 | [50.91,55.70] | 53.05 | [48.20,58.10] | 44 |
|  |  | m | [43.70,86.00] | 56.88 | 7.87 | [54.54,59.21] | 56.70 | [51.93,59.33] | 46 |
|  | Southern Highlands | f | [34.60,80.90] | 50.69 | 8.96 | [47.94,53.45] | 49.90 | [44.60,56.85] | 43 |
|  |  | m | [41.20,67.80] | 54.44 | 5.91 | [52.64,56.24] | 53.95 | [49.75,58.20] | 44 |
|  | West Coast | f | [37.30,83.30] | 53.91 | 10.67 | [50.66,57.15] | 53.05 | [46.55,57.30] | 44 |
|  |  | m | [46.30,87.40] | 58.79 | 7.59 | [56.46,61.13] | 57.50 | [54.25,62.85] | 43 |
| Body fat percentage (%) | Central Highlands | f | [9.80,30.80] | 20.41 | 4.55 | [19.02,21.79] | 20.85 | [17.10,22.88] | 44 |
|  |  | m | [6.90,23.40] | 13.30 | 3.36 | [12.30,14.30] | 12.70 | [11.38,14.80] | 46 |
|  | Southern Highlands | f | [11.20,39.20] | 19.02 | 5.21 | [17.42,20.62] | 18.30 | [15.70,21.35] | 43 |
|  |  | m | [6.00,17.60] | 12.26 | 2.56 | [11.48,13.03] | 12.10 | [10.88,13.30] | 44 |
|  | West Coast | f | [8.60,77.60] | 22.17 | 10.61 | [18.94,25.40] | 20.30 | [16.48,25.00] | 44 |
|  |  | m | [6.40,26.60] | 13.74 | 3.70 | [12.60,14.88] | 13.10 | [11.20,16.15] | 43 |
| Body mass index (kg / m<sup>2</sup>) | Central Highlands | f | [17.60,29.40] | 22.66 | 2.88 | [21.79,23.54] | 22.95 | [20.35,24.53] | 44 |
|  |  | m | [11.90,27.70] | 20.79 | 2.51 | [20.05,21.54] | 20.60 | [19.70,21.95] | 46 |
|  | Southern Highlands | f | [16.40,35.00] | 21.61 | 3.45 | [20.54,22.67] | 21.10 | [19.40,23.15] | 43 |
|  |  | m | [16.20,23.90] | 20.19 | 1.66 | [19.68,20.69] | 20.10 | [19.20,21.00] | 44 |
|  | West Coast | f | [13.30,31.70] | 22.57 | 4.40 | [21.23,23.91] | 22.00 | [19.83,25.40] | 44 |
|  |  | m | [16.50,29.80] | 21.35 | 2.43 | [20.60,22.10] | 20.90 | [19.65,22.95] | 43 |
| Waist circumference (cm) | Central Highlands | f | [62.70,94.80] | 76.73 | 8.14 | [74.26,79.21] | 74.85 | [70.88,82.48] | 44 |
|  |  | m | [63.50,99.00] | 73.79 | 6.93 | [71.73,75.85] | 72.75 | [69.20,76.60] | 46 |
|  | Southern Highlands | f | [57.50,108.00] | 75.13 | 9.37 | [72.25,78.01] | 75.00 | [69.50,79.25] | 43 |
|  |  | m | [65.00,85.00] | 72.57 | 4.07 | [71.34,73.81] | 72.35 | [70.00,74.13] | 44 |
|  | West Coast | f | [57.00,103.00] | 77.02 | 10.91 | [73.71,80.34] | 75.50 | [70.63,82.25] | 44 |
|  |  | m | [62.00,105.50] | 74.37 | 6.80 | [72.28,76.46] | 74.00 | [70.75,75.50] | 43 |
| Hip circumference (cm) | Central Highlands | f | [76.00,104.00] | 89.31 | 6.77 | [87.25,91.36] | 88.70 | [84.68,93.93] | 44 |
|  |  | m | [77.50,107.00] | 86.56 | 5.07 | [85.05,88.07] | 85.80 | [84.58,88.28] | 46 |
|  | Southern Highlands | f | [78.50,109.00] | 89.13 | 6.30 | [87.19,91.07] | 88.00 | [84.25,93.75] | 43 |
|  |  | m | [76.00,92.50] | 84.99 | 3.96 | [83.78,86.19] | 84.65 | [82.75,88.00] | 44 |
|  | West Coast | f | [75.00,110.00] | 91.36 | 7.68 | [89.03,93.70] | 91.00 | [87.75,95.00] | 44 |
|  |  | m | [77.00,107.00] | 86.06 | 5.33 | [84.42,87.70] | 85.00 | [82.50,89.00] | 43 |
| Heart pulse (beats / min) | Central Highlands | f | [62.00,108.00] | 80.95 | 12.25 | [77.23,84.68] | 80.00 | [71.00,89.25] | 44 |
|  |  | m | [58.00,99.00] | 77.76 | 12.16 | [74.15,81.37] | 76.50 | [68.25,88.00] | 46 |
|  | Southern Highlands | f | [33.00,123.00] | 78.79 | 16.31 | [73.77,83.81] | 75.00 | [68.00,89.50] | 43 |
|  |  | m | [50.00,99.00] | 72.25 | 12.95 | [68.31,76.19] | 70.00 | [64.75,79.50] | 44 |
|  | West Coast | f | [63.00,117.00] | 83.91 | 13.87 | [79.64,88.17] | 83.00 | [74.50,89.50] | 43 |
|  |  | m | [47.00,93.00] | 68.58 | 10.58 | [65.32,71.84] | 69.00 | [61.50,74.50] | 43 |
| Basal metabolic rate (cal / day) | Central Highlands | f | [812.00,1621.00] | 1,203.05 | 168.83 | [1151.72,1254.37] | 1,196.00 | [1089.00,1293.75] | 44 |
|  |  | m | [948.00,1918.00] | 1,306.74 | 193.14 | [1249.38,1364.10] | 1,281.50 | [1162.75,1403.50] | 46 |
|  | Southern Highlands | f | [751.00,1675.00] | 1,123.19 | 196.42 | [1062.74,1183.64] | 1,126.00 | [953.00,1292.50] | 43 |
|  |  | m | [886.00,1546.00] | 1,254.57 | 150.52 | [1208.81,1300.33] | 1,247.50 | [1160.00,1339.75] | 44 |
|  | West Coast | f | [772.00,1966.00] | 1,207.57 | 247.87 | [1132.21,1282.93] | 1,162.00 | [1079.50,1327.25] | 44 |
|  |  | m | [995.00,1879.00] | 1,330.07 | 176.16 | [1275.86,1384.28] | 1,305.00 | [1214.00,1461.00] | 43 |
| Blood pressure - Systolic (mmHg) | Central Highlands | f | [100.00,149.00] | 126.84 | 12.91 | [122.92,130.76] | 129.00 | [117.75,135.00] | 44 |
|  |  | m | [112.00,147.00] | 127.22 | 8.51 | [124.69,129.74] | 127.50 | [122.00,133.75] | 46 |
|  | Southern Highlands | f | [96.00,162.00] | 131.91 | 15.71 | [127.07,136.74] | 132.00 | [122.00,142.50] | 43 |
|  |  | m | [112.00,157.00] | 131.18 | 10.51 | [127.99,134.38] | 131.00 | [123.75,138.00] | 44 |
|  | West Coast | f | [94.00,171.00] | 120.12 | 19.33 | [114.17,126.06] | 113.00 | [106.50,129.00] | 43 |
|  |  | m | [94.00,155.00] | 122.81 | 14.30 | [118.41,127.21] | 121.00 | [113.50,131.50] | 43 |
| Blood pressure - Diastolic (mmHg) | Central Highlands | f | [60.00,99.00] | 80.80 | 10.14 | [77.71,83.88] | 81.50 | [73.00,88.00] | 44 |
|  |  | m | [60.00,94.00] | 76.22 | 9.33 | [73.45,78.99] | 76.00 | [70.00,83.00] | 46 |
|  | Southern Highlands | f | [41.00,114.00] | 82.40 | 14.29 | [78.00,86.79] | 83.00 | [77.00,92.00] | 43 |
|  |  | m | [62.00,109.00] | 83.93 | 11.79 | [80.35,87.52] | 83.50 | [74.00,93.00] | 44 |
|  | West Coast | f | [51.00,121.00] | 77.28 | 15.20 | [72.60,81.96] | 73.00 | [67.00,81.50] | 43 |
|  |  | m | [53.00,109.00] | 74.16 | 12.85 | [70.21,78.12] | 74.00 | [66.00,79.50] | 43 |

**Supplemental Table 2.**
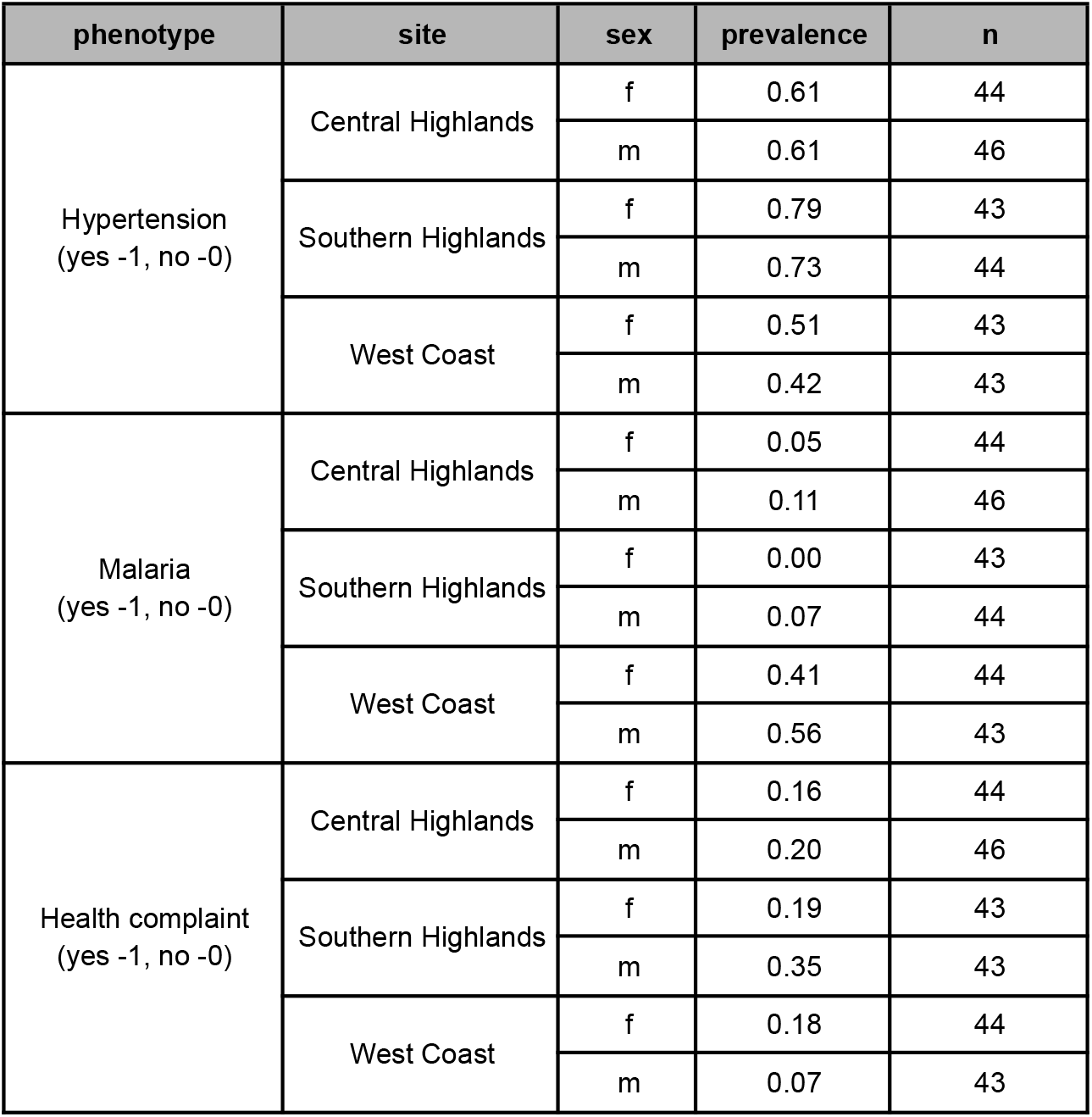
Descriptive statistics for binary phenotypes collected stratified by site and community. Prevalence (proportion of participants responding “yes”) and total number of samples with a value for each phenotype are listed.

**Supplemental Table 3.**
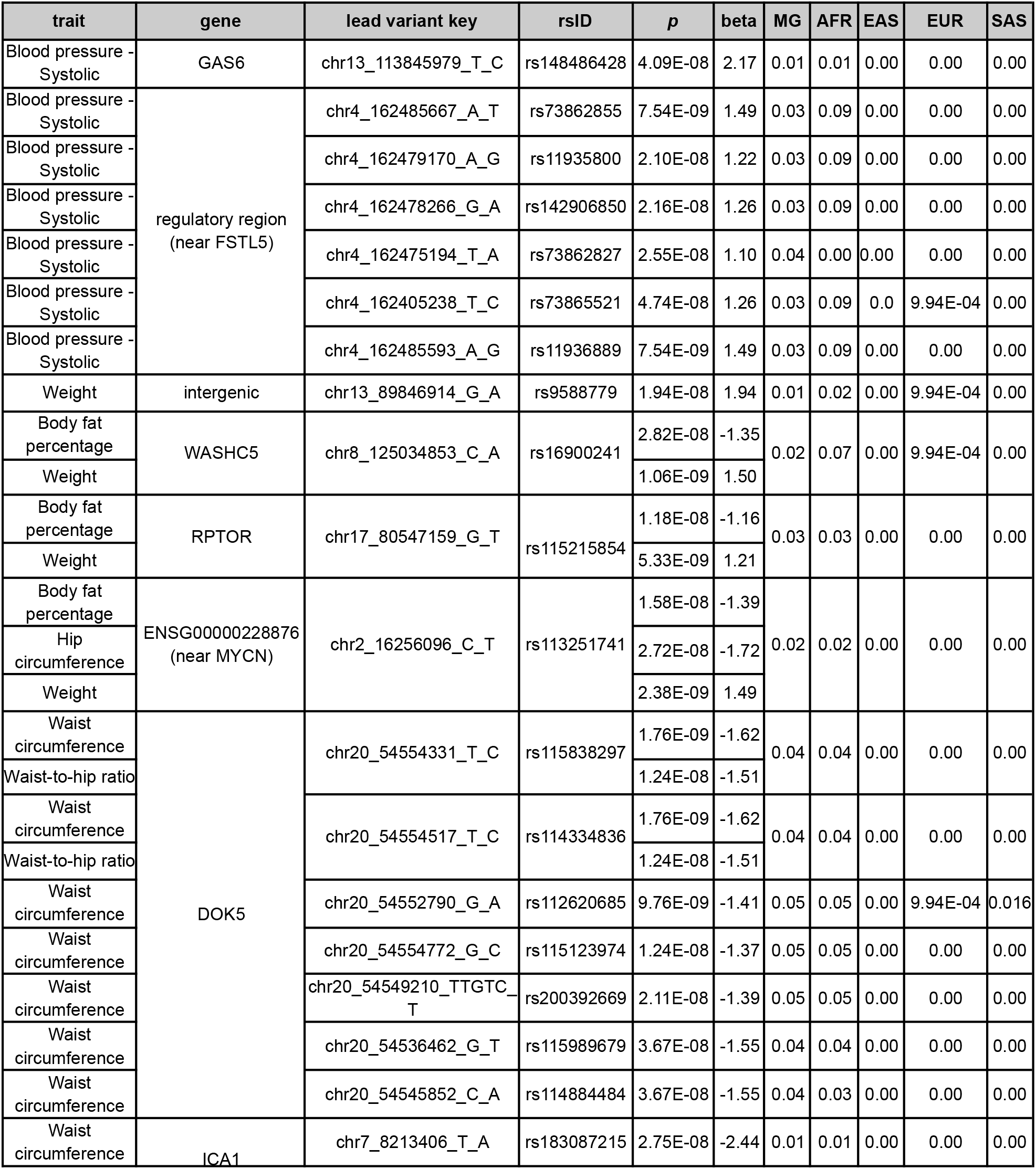
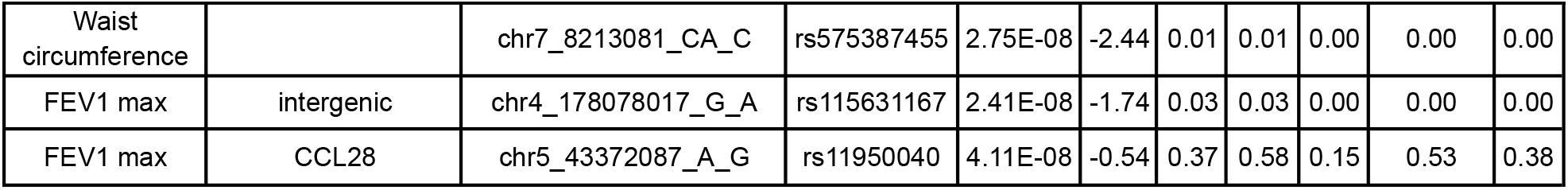
Summary statistics for all 22 variants with at least one significant GWAS association. The *p*-values and beta coefficients for each variant and trait associated are listed. The associated variant key (chr_pos_ref_alt), rsID, VEP annotated gene (or closest coding gene in parenthesis), and allele frequencies are included (MG=Malagasy cohort from this study, AFR=1000 Genomes African populations, EAS=1000 Genomes East Asian populations, EUR=1000 Genomes European populations, SAS=1000 Genomes South Asian populations).

**Supplemental Table 4.** Summary of all 16 GWAS run in this study. For each phenotype, the lambda GC, number of hits, number of independent hits as determined by conditional analysis, the number of loci, sample size, and the number of cases and controls if applicable.

| Phenotype Name | Lambda GC | Hits | Independent Hits | Loci | Sample Size | Cases | Controls |
| --- | --- | --- | --- | --- | --- | --- | --- |
| Basal metabolic rate | 1.0324 | 0 | 0 | 0 | 214 | 0 | 0 |
| Body mass index | 1.0495 | 0 | 0 | 0 | 214 | 0 | 0 |
| Blood pressure - Diastolic | 0.9925 | 0 | 0 | 0 | 213 | 0 | 0 |
| Blood pressure - Systolic | 1.0182 | 7 | 3 | 2 | 213 | 0 | 0 |
| FEV1 max | 1.0395 | 2 | 2 | 2 | 214 | 0 | 0 |
| FVC max | 1.0047 | 0 | 0 | 0 | 214 | 0 | 0 |
| Heart pulse | 1.0005 | 0 | 0 | 0 | 213 | 0 | 0 |
| Height | 1.0269 | 0 | 0 | 0 | 214 | 0 | 0 |
| Hip circumference | 1.0469 | 1 | 1 | 1 | 214 | 0 | 0 |
| Hypertension | 1.1136 | 0 | 0 | 0 | 213 | 129 | 84 |
| Lean body mass | 1.0394 | 0 | 0 | 0 | 214 | 0 | 0 |
| Malaria | 1.2579 | 0 | 0 | 0 | 214 | 46 | 168 |
| Body fat percentage | 1.0451 | 3 | 4 | 3 | 214 | 0 | 0 |
| Waist circumference | 1.0422 | 9 | 2 | 2 | 214 | 0 | 0 |
| Weight | 1.0402 | 4 | 5 | 4 | 214 | 0 | 0 |

**Supplemental Figure 1.**
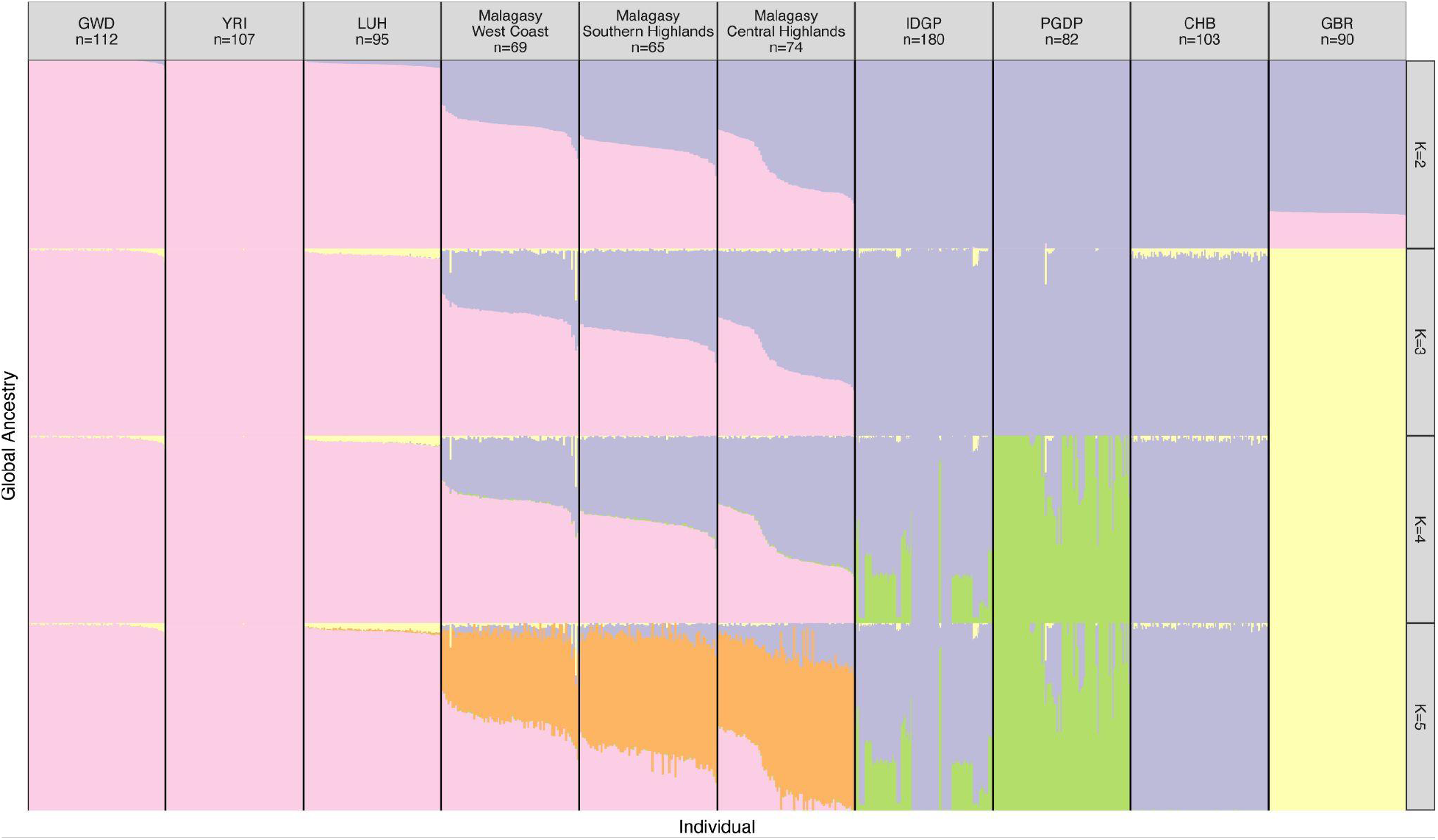
ADMIXTURE plot for K=2 to K=5, including Malagasy cohort (West Coast, Central Highlands, Southern Highlands) and relevant reference populations from 1000 Genomes (GWD, YRI, LUH CHB, GBR) and additional reference samples from IDGP and PGDP. Global ancestry bars are colored by proportion of ancestry from each inferred source (2-5 sources depending on K value).

**Supplemental Figure 2.**
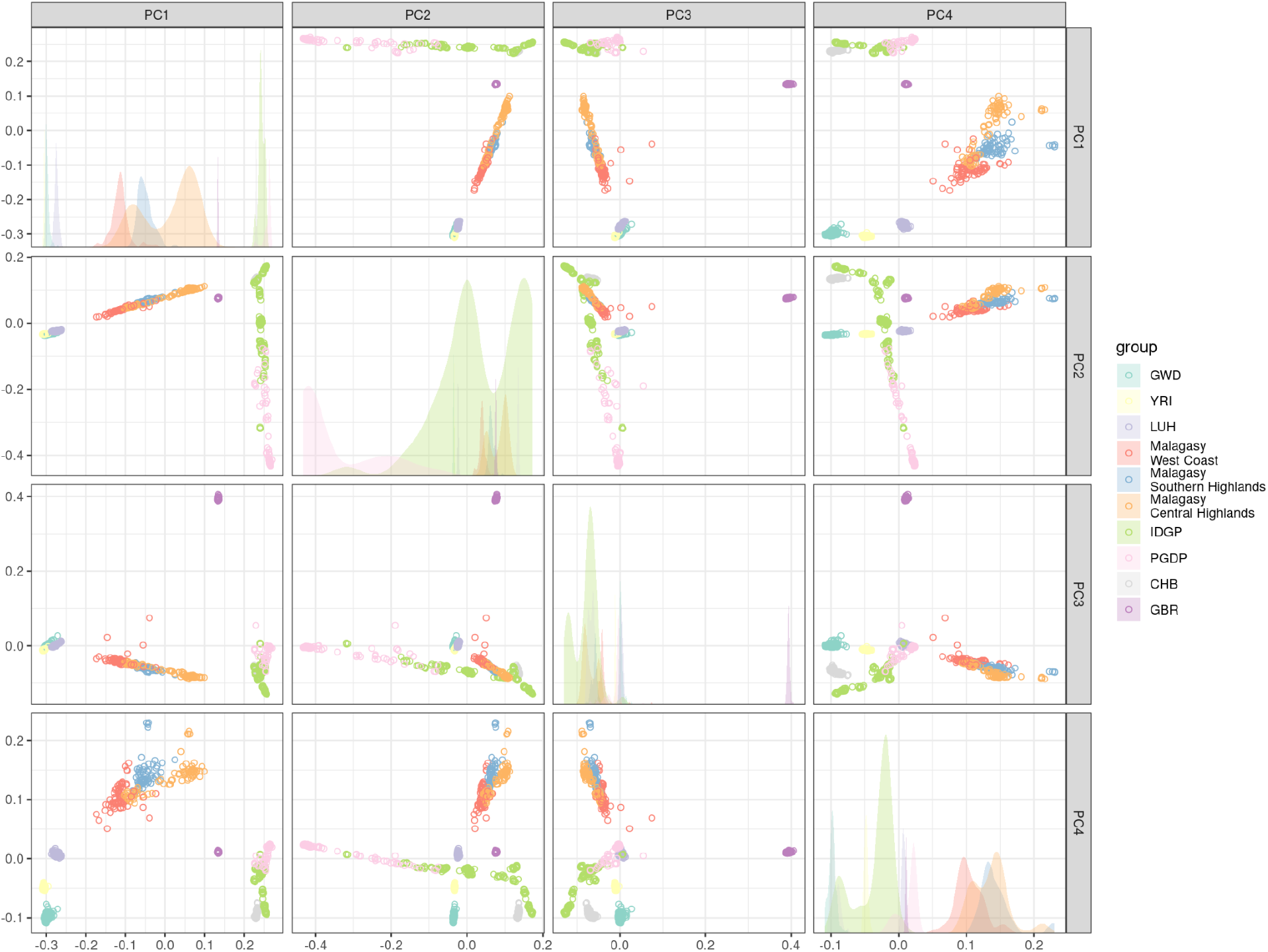
Grid of PCs 1-4 for Malagasy cohort (West Coast, Central Highlands, Southern Highlands) and relevant reference populations from 1000 Genomes (GWD, YRI, LUH CHB, GBR) and additional reference samples from IDGP and PGDP. Diagonal shows density distributions across the X axis for each PC. Points in scatterplot or density fills are colored by group labels.

**Supplemental Figure 3.**
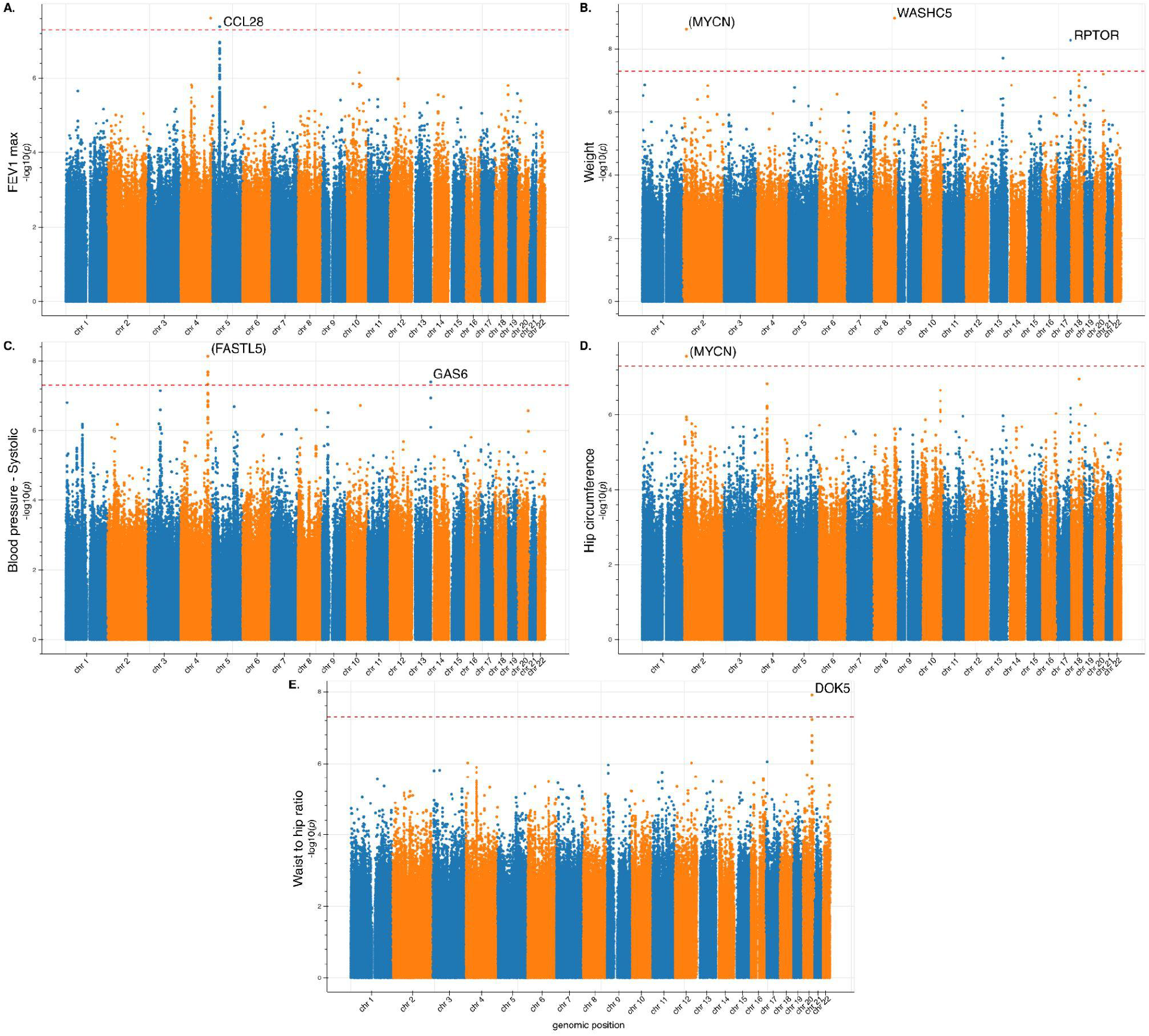
GWAS Manhattan plots for traits with significant associations. *P*-values for all autosomal variants passing GWAS QC on a −log10 scale for **A)** FEV1 max, **B)** weight, **C)** systolic blood pressure, **D)** hip circumference, and **E)** waist to hip ratio. Annotated gene for top variant, or nearest protein coding gene in parenthesis, is listed for each locus reaching genome-wide significance (*p* < 5e-8, red dashed line). Lead variants without an annotated gene are intergenic.

**Supplemental Figure 4.**
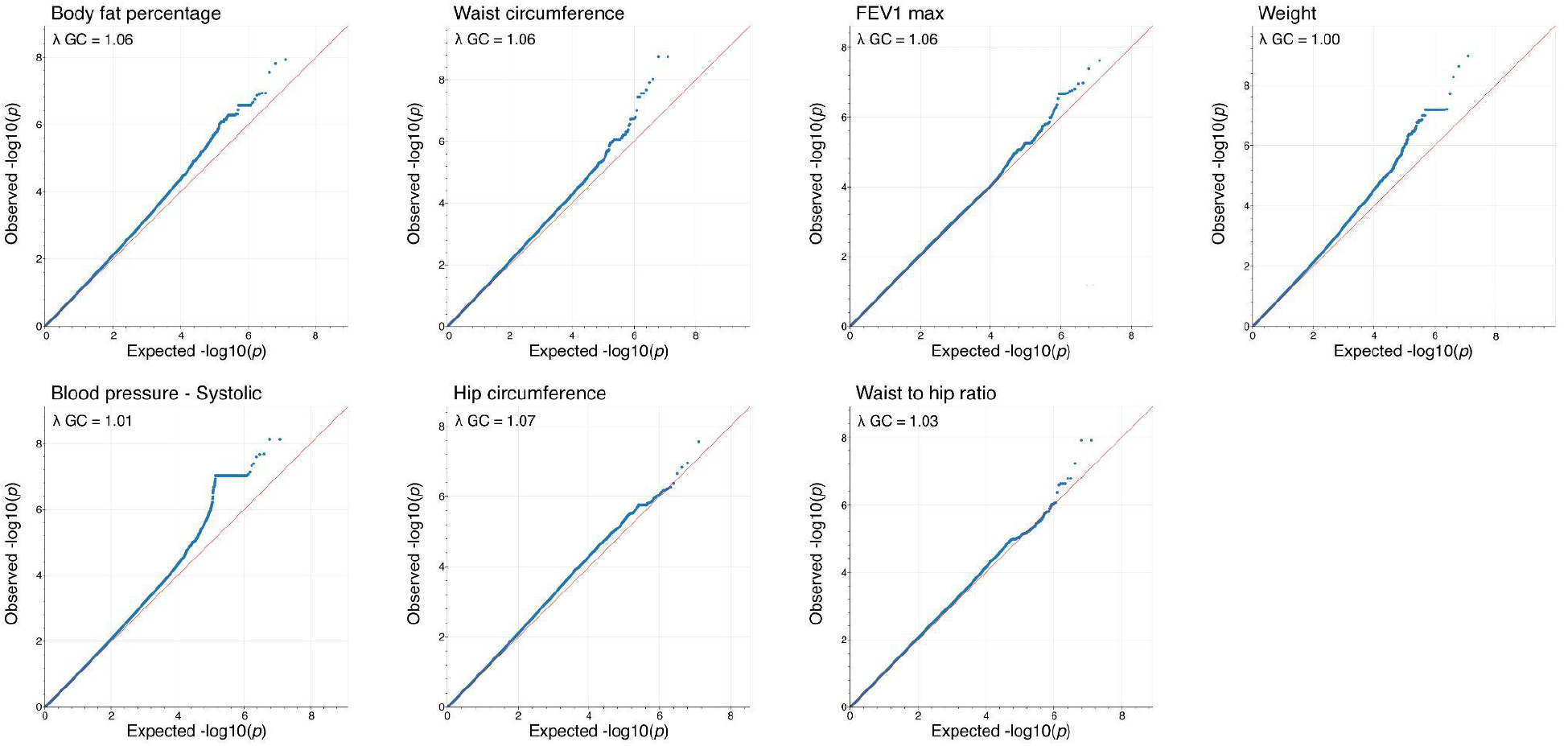
Quantile-quantile plots for traits with significant associations. Observed vs expected *p*-values for all autosomal variants passing GWAS QC on a −log10 scale for body fat percentage, waist circumference, FEV1 max, weight, systolic blood pressure, hip circumference, and waist to hip ratio. Genomic inflation factor (λ GC) for each trait is annotated for each QQ plot.

